# Translating multi-omics complexity into sparse prognostic biomarkers for multiple myeloma

**DOI:** 10.64898/2026.08.21.746155

**Authors:** Benedikt Obermayer, Manuela Benary, Jan Krönke, Philipp Mertins, Dieter Beule

## Abstract

Multiple myeloma (MM) exhibits profound molecular heterogeneity, yet current risk stratification relies on cytogenetics or single-omics signatures that often fail to capture cross-layer regulatory complexity. We re-analyzed a multi-omics dataset integrating copy-number, transcriptomic, proteomic, and phosphoproteomic data to dissect how common genomic driver alterations propagate through the molecular cascade. Supervised classification demonstrated that downstream layers, particularly the proteome and phosphoproteome, classify genomic events more accurately than primary genomic or transcriptomic data. Intriguingly, trans-acting features alone were sufficient for classification, indicating that while direct dosage effects manifest at the RNA level, downstream network responses dominate the proteomic state.

Multi-omics factor analysis (MOFA2) identified a continuous latent axis predicting progression-free and overall survival independent of R-ISS. This factor captured a gain(1q)/del(13q) axis modulated by immune infiltration and NSD2 expression, integrating variance across all four modalities. To enable clinical translation, we derived sparse, single-modality proxies using elastic net regression. An RNA proxy faithfully recapitulated the multi-omic factor and validated independently in published microarray and RNAseq cohorts, demonstrating robust prognostic utility across treatment eras. These findings reveal that multi-omics integration uncovers hidden prognostic axes obscured by single-omics analyses, and that sparse proxies can bridge the gap between complex discovery and clinical implementation.

## Introduction

Multiple myeloma is a genetically and biologically heterogeneous plasma cell malignancy (van de Donk et al., 2021), and prognosis remains difficult to capture with current clinical and genomic risk models. Risk stratification remains anchored in clinical staging and recurrent cytogenetic abnormalities such as deletion of 17p (del(17p)), gain of chromosome arm 1q (gain(1q)), and the IGH translocations t(4;14) and t(14;16) (Avet-Loiseau et al., 2012; Santra et al., 2003), with the revised International Staging System providing an important but incomplete framework for classification (Palumbo et al., 2015; Rajkumar, 2024). Recent large-scale whole-genome sequencing has expanded the catalog of coding and non-coding drivers in myeloma and its precursors, and genomic scores derived from these features show promise for stratifying progression risk beyond clinical staging (Alberge et al., 2025; Chapman et al., 2011). On the other hand, transcriptome-based risk scores have also added value in myeloma, with the GEP-70 signature one of the best-established examples of prognostic expression profiling in this disease (Heuck et al., 2014; Kumar et al., 2011). Even so, transcript abundance is only an indirect readout of tumor state, and mRNA levels often diverge from protein abundance and signaling activity in cancer, leaving important biology invisible to transcript-only models (Chen et al., 2002; Kochavi et al., 2023).

This limitation motivates studies that combine data across multiple molecular layers to uncover coordinated regulatory programs that are not apparent from a single assay and thus separate direct genomic effects from broader downstream consequences (Mani et al., 2022; Ortiz-Estevez et al., 2021). Ramberger et al. recently showed that the molecular landscape of multiple myeloma is shaped not only by chromosomal alterations but also by post-transcriptional regulation, and they identified a prognostic protein signature associated with aggressive disease independent of established risk factors (Ramberger et al., 2024). These findings support the idea that clinically relevant risk states are distributed across molecular processes and cannot easily be captured by one assay alone.

A relevant biological question is therefore how recurrent genomic lesions propagate through the tumor system. Some alterations act locally, but others exert trans effects that transmit the consequences of a given lesion into downstream pathways. For instance, gain(1q) is associated with altered proliferation, apoptosis resistance, metabolic reprogramming, drug resistance, and immune evasion (Kuroki et al., 2025; Schmidt et al., 2021). The importance of context is reinforced by emerging spatial work showing that myeloma biology is niche-dependent, with distinct microenvironmental programs in bone marrow and extramedullary disease that can themselves stratify outcome (Ohlstrom et al., 2026).

These observations raise a translational challenge. Latent molecular programs are easy to discover in unsupervised analysis, but they are not immediately useful in the clinic unless they can be reduced to simple, robust, and assayable biomarkers. Sparse linear models are well suited to this task because they can convert complex multi-omic signals into compact proxies that preserve predictive information and can be validated in independent cohorts (Argelaguet et al., 2020; Basu et al., 2018; Min et al., 2018; Traeuble et al., 2026).

Here, we reanalyze the Ramberger et al. proteogenomic cohort (Ramberger et al., 2024) and integrate copy-number, transcriptomic, proteomic, and phosphoproteomic data to ask how genomic events propagate across molecular layers, which latent factors carry prognostic information, and whether those latent signals can be recapitulated by sparse single-modality proxies. In doing so, we aim to connect discovery-level proteogenomic biology with biomarkers that are simple enough to support clinical translation while retaining the prognostic power of the broader multi-omic state.

## Results

### Multi-omic characterization of multiple myeloma

From the initial cohort of 138 primary patient-derived plasma cell malignancies, we retained 120 patients with complete multi-omic data, of which 104 were newly diagnosed with multiple myeloma, 13 had plasmacytoid leukemia and 3 had monoclonal gammopathy of undetermined significance (MGUS). Primary genomic alterations (a hyperdiploid karyotype (HRD) with amplifications of odd-numbered chromosomes, or translocations involving the immunoglobulin heavy chain (*IgH*) enhancer on chromosome 14) and secondary events (gains or losses on chromosomes 1, 9, 13 or 17), derived from fluorescence in-situ hybridization (FISH), are summarized in **Figure 1A**. Further modality-specific feature filtering yielded 626 loci assessed by shallow nanopore whole-genome DNA sequencing for copy-number status (“CNV”), 5477 genes profiled by bulk RNA sequencing (“RNA”), as well as TMT-based quantification of proteome (“protein”, n=6383) and phosphoproteome (“phospho”, n=6446) (**Figure 1B**). We used batch correction to harmonize samples with varying RNA quality and CD138 enrichment strategy (**Figure S1**).

**Figure 1:**
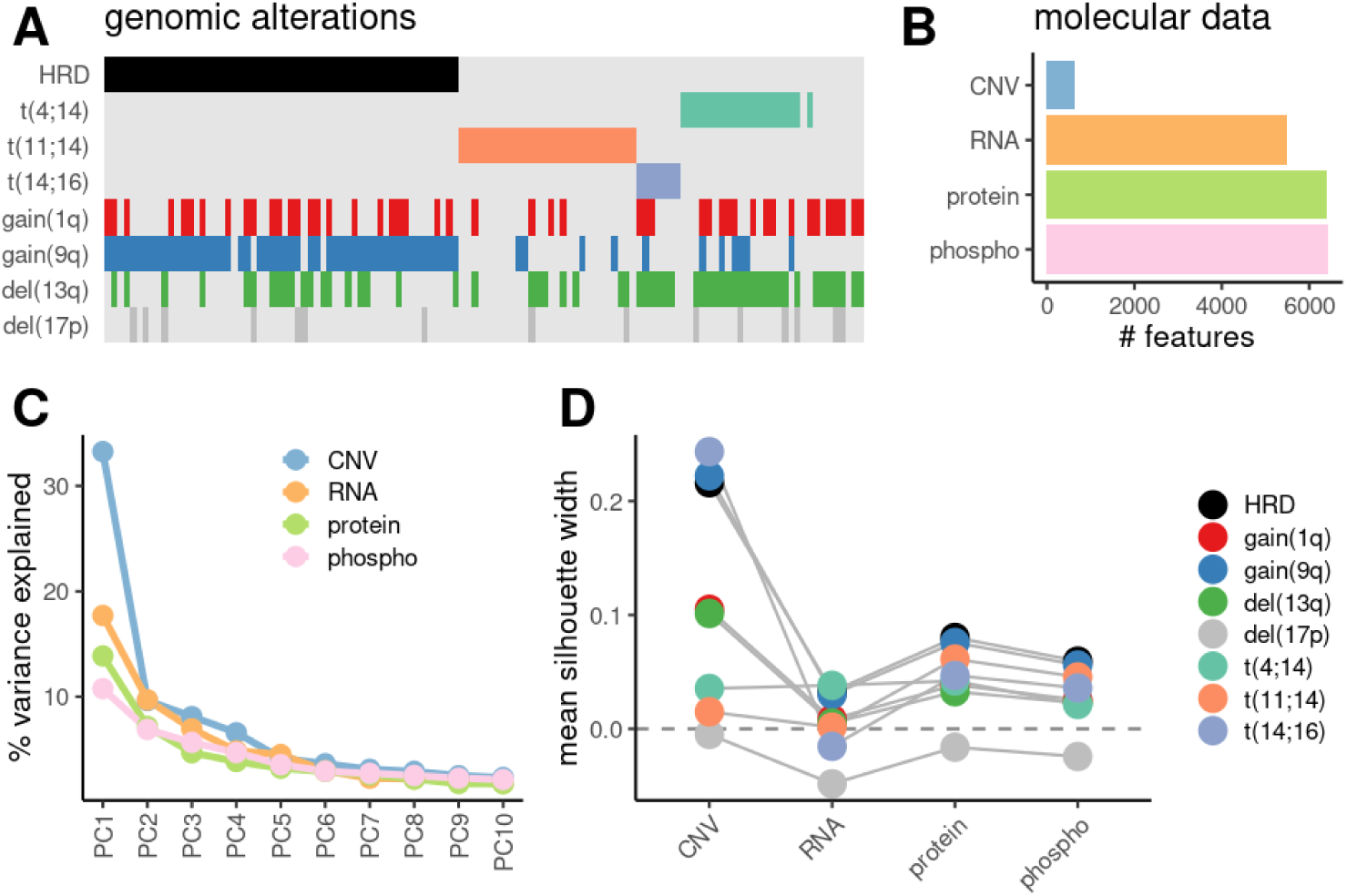
Multi-omic characterization of multiple myeloma. **A**: Overview of primary and secondary genomic driver events. **B**: number of features profiled per molecular modality after QC filtering (CNV: shallow nanopore whole-genome sequencing; RNA: bulk RNAseq; protein: proteomics; phospho: phosphoproteomics). **C**: Percentage of variance explained per principal component in each modality. **D**: unsupervised power to detect presence of genomic alterations measured by the mean silhouette width in PC space for each modality.

A principal component (PC) analysis for each modality showed a substantial amount of variance concentrated in the first few components, particularly for the CNV and RNA modalities (**Figure 1C**). Measuring the separation of samples with and without a particular genomic alteration in that PC space using silhouette analysis indicated that some CNV-defined events (HRD, gain(9q), gain(1q) and del(13q)) produce a strong clustering in CNV space, while the translocations t(4;14) and t(11;14) and all events in the other modalities do not show strong separation in such an unsupervised analysis.

### Molecular signatures of genomic driver events

To characterize the molecular consequences of different genomic alterations and to identify information-rich features in each modality, we used partial least-squares discriminant analysis (PLS-DA), which in contrast to the principal components analysis before is a supervised dimensionality reduction method that maximizes covariance with a target variable (Rohart et al., 2017). Using gene set enrichment analysis (Zyla et al., 2019), we found that this approach prioritized genes in expected pathways, such as terms relating to nonsense-mediated decay or translation for del(13q) due to loss of DIS3 expression (Skerget et al., 2024), metabolic reprogramming for HRD and gain(9q), MYC targets for t(11;14) and cell-cycle and p53-related genes for del(17p) (**Figure S2A**). The resulting separation of samples with and without a specific alteration, quantified using the area-under-curve metric (AUC), is shown in **Figure 2A** for a dense PLS-DA (using all features of each modality) as well as a sparse version (prioritizing at most 150 features in each modality). Average AUC values reach 0.85-0.92 for RNA, protein and phospho for all genomic drivers except del(17p), which is a rare and heterogeneous event (partial vs. biallelic loss of TP53) that cannot be confidently assessed in a cross-validation setting. As expected, the CNV modality is not informative for translocations. The sparse methods tend to outperform the dense ones, indicating that the molecular signatures of these alterations coalesce around a limited set of highly informative features.

**Figure 2:**
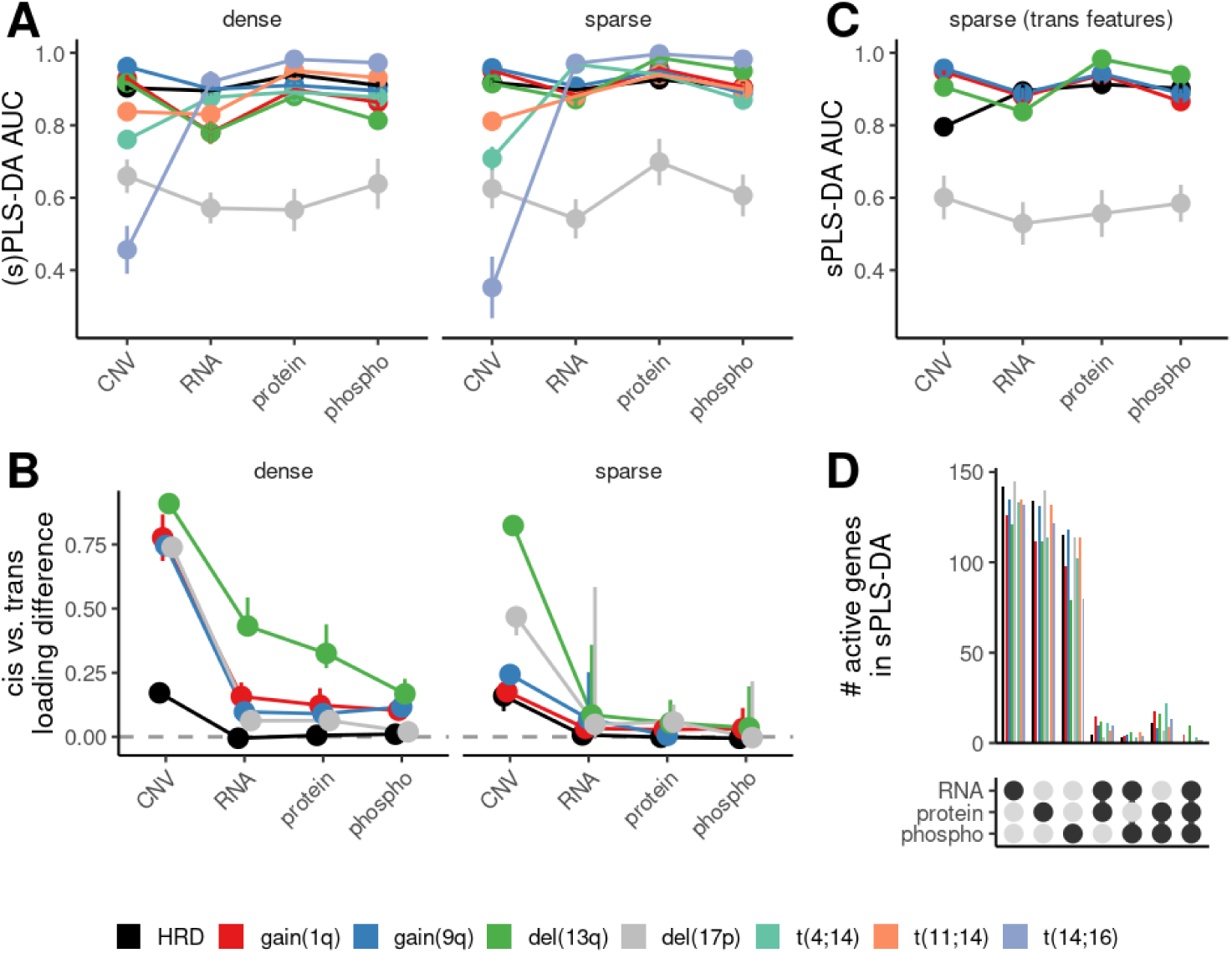
Molecular signatures of genomic driver events characterized by partial least squares discriminant analysis (PLS-DA). **A**: area-under-curve (AUC) values from 5fold cross-validation of PLS-DA using all features (“dense”) or 150 features (“sparse”). Error bars denote standard deviation. **B**: Difference in median squared loading between cis or trans features for the (s)PLS-DA models in **A**. Error bars denote 95% confidence intervals from bootstrapping. **C**: AUC values as in **A** but for sPLS-DA using only trans features. **D**: Bar plot showing the number of active genes in sPLS-DA models in **A** shared between different modalities.

We next asked whether relevant features with high loadings were concentrated in the affected genomic loci (“*cis*”) or elsewhere in the genome (“*trans*”) and quantified differences in the loading distributions (**Figure S2B**) using the difference of the median squared loading (**Figure 2B**). Consistent with a central dogma-like flow of information from DNA via RNA to protein and post-translational signaling, the upstream layers showed stronger localization in *cis*, while the downstream layers propagated the signal towards (genomically) distant nodes in the regulatory network. Strikingly, this effect was much less pronounced for the sparse models, suggesting that the most informative features are not primarily located in *cis*. In fact, sparse models using only *trans* features performed equally well (**Figure 2C**). Finally, we investigated the degree of coordination between the layers and found very few active genes shared between models using different modalities for the same target (**Figure 2D**). Together with the limited overlap of enriched pathways seen in **Figure S2A**, this emphasizes the notion that each modality profiles different and non-redundant facets of the molecular consequences of each alteration.

### Multi-omic factor analysis

To further investigate variance shared between or specific to molecular modalities, we continued with an unsupervised analysis of the dataset using MOFA2 (Argelaguet et al., 2020). Among 10 factors chosen initially, we saw four with substantial contributions from all modalities (F1, F3, F5 and F6), while the others were restricted to specific layers (**Figure 3A**).

**Figure 3:**
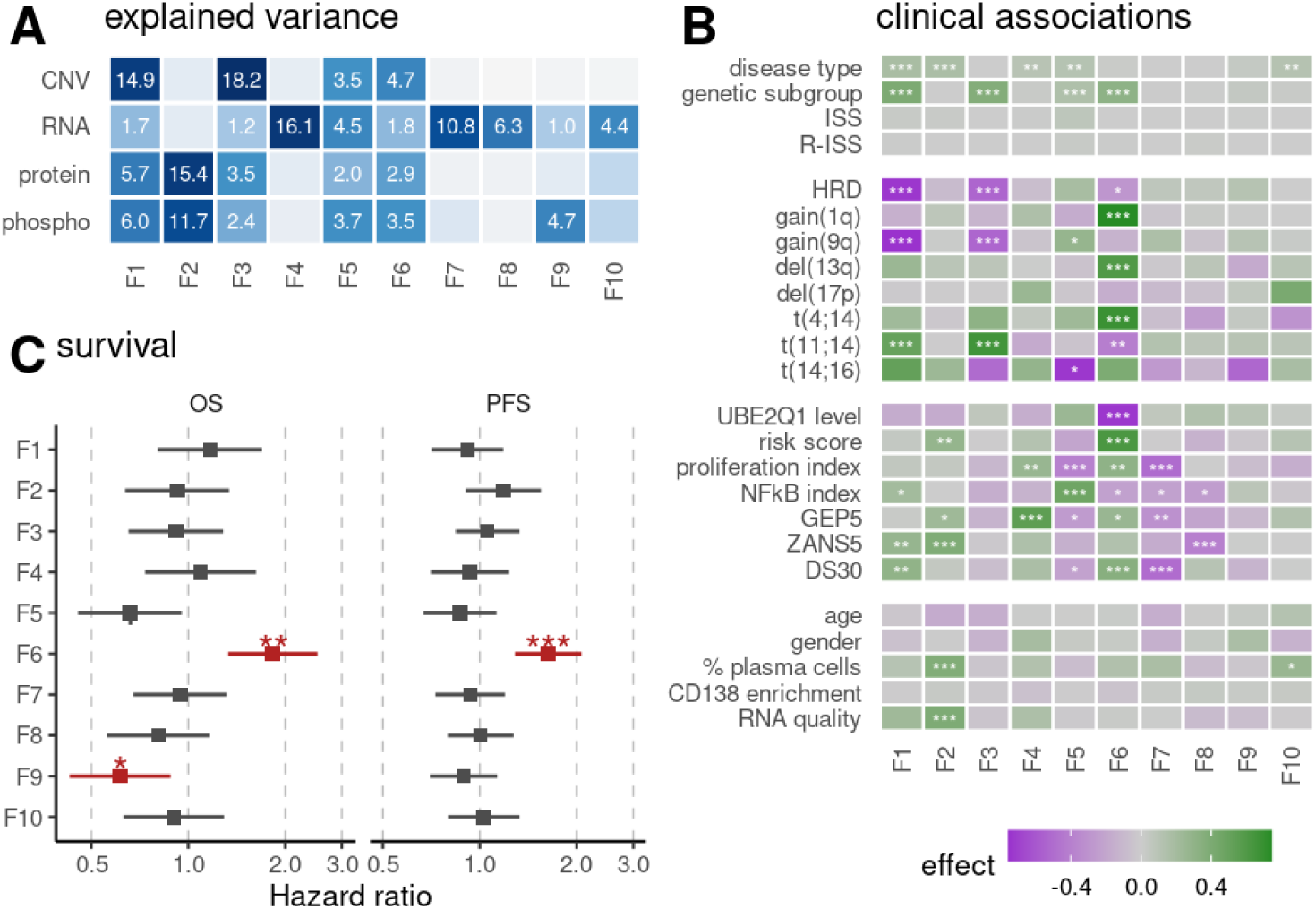
Unsupervised multi-omic factor analysis using MOFA2. **A**: percentage of variance explained by each factor per modality. **B**: associations between factor loadings and different clinical and molecular variables. P-values and effect sizes result from Wilcoxon tests and rank-biserial correlations for binary variables, from Kruskal-Wallis tests and the η^2^ statistics for categorical variables, and as Spearman correlation for numerical variables. **C**: Hazard ratios for overall (OS) and progression-free (PFS) survival estimated from Cox regression analysis of MOFA2 factors using R-ISS as covariate. P-values in **B+C** adjusted for multiple comparisons using the BH method. ***: p < 0.001, **: p < 0.01, *: p < 0.05.

Consistent with this observation, we observed strong associations of these four factors with most of the genomic alterations, and disease type or (primary) genetic subgroup generally (**Figure 3B**). In particular, F6 was strongly associated with gain(1q) and del(13q), which are known as detrimental independently and (even more so) in combination (Skerget et al., 2024), but also with the translocations t(4;14) and t(11;14), and finally with the level of UBE2Q1 (Chang et al., 2015), the protein-based risk score identified in the original study by Ramberger et al, and a recently proposed 30-gene dissemination score (Bohra et al., 2026). In contrast, various scores derived in the literature from transcriptome data, such as a proliferation score (Bergsagel et al., 2005), an NF-κB index (Ang et al., 2024), a 5-gene version of the GEP-70 signature (Heuck et al., 2014) and another “ZANS5” 5-gene signature developed by Zamani-Ahmadmahmudi, Nassiri and Soltaninezhad (Zamani-Ahmadmahmudi et al., 2021) were associated with other factors, including some that explained variance only in the RNA modality. None of the factors were associated with ISS, R-ISS, age, gender or CD138 enrichment strategy, and other technical factors such as RNA quality and plasma cell content were restricted to F2. Interestingly, the strongest signals in a gene set enrichment analysis (**Figure S3**) showed up in modality-specific factors, suggesting that these factors are more easily interpreted with this technique, while the factors shared across modalities integrate various biological signals.

### Prognostic value of factor 6

Next, we tested all factors for prognostic power beyond current clinical practice using Cox regression with R-ISS as covariate (**Figure 3C**), which revealed F6 as the only factor with prognostic value both in overall survival and progression-free survival, in agreement with the strong associations seen in **Figure 3B**. Indeed, Kaplan-Meier curves showed a significant difference in survival between patients with high or low F6 score (**Figure 4A+B**). Inspecting F6 feature loadings (**Figure 4C**), we recovered signatures of the genomic events found in **Figure 3B**, such as gain(1q) and del(13q) in the CNV modality, and the overexpression of NSD2 caused by t(4;14) in the RNA and protein layer (Santra et al., 2003), but also an immune-infiltration signature through genes like SELPLG, LSP1, LAPTM5, LAMP3 and EPSTI1 (Karlsson et al., 2021). To test our initial choice of parameters, we further ran MOFA2 with different numbers of factors and found that in all cases with at least 10 factors (which is roughly where the explained variance per modality starts to saturate) exactly one of them is highly correlated to F6, which then is also the only factor with prognostic value, indicating that the multi-omic axis captured by F6 can be robustly isolated. We obtained comparable results when running MOFA2 only on data from multiple myeloma patients (i.e., without MGUS or PCL) or only on CD138+ input (**Figure S4**).

**Figure 4:**
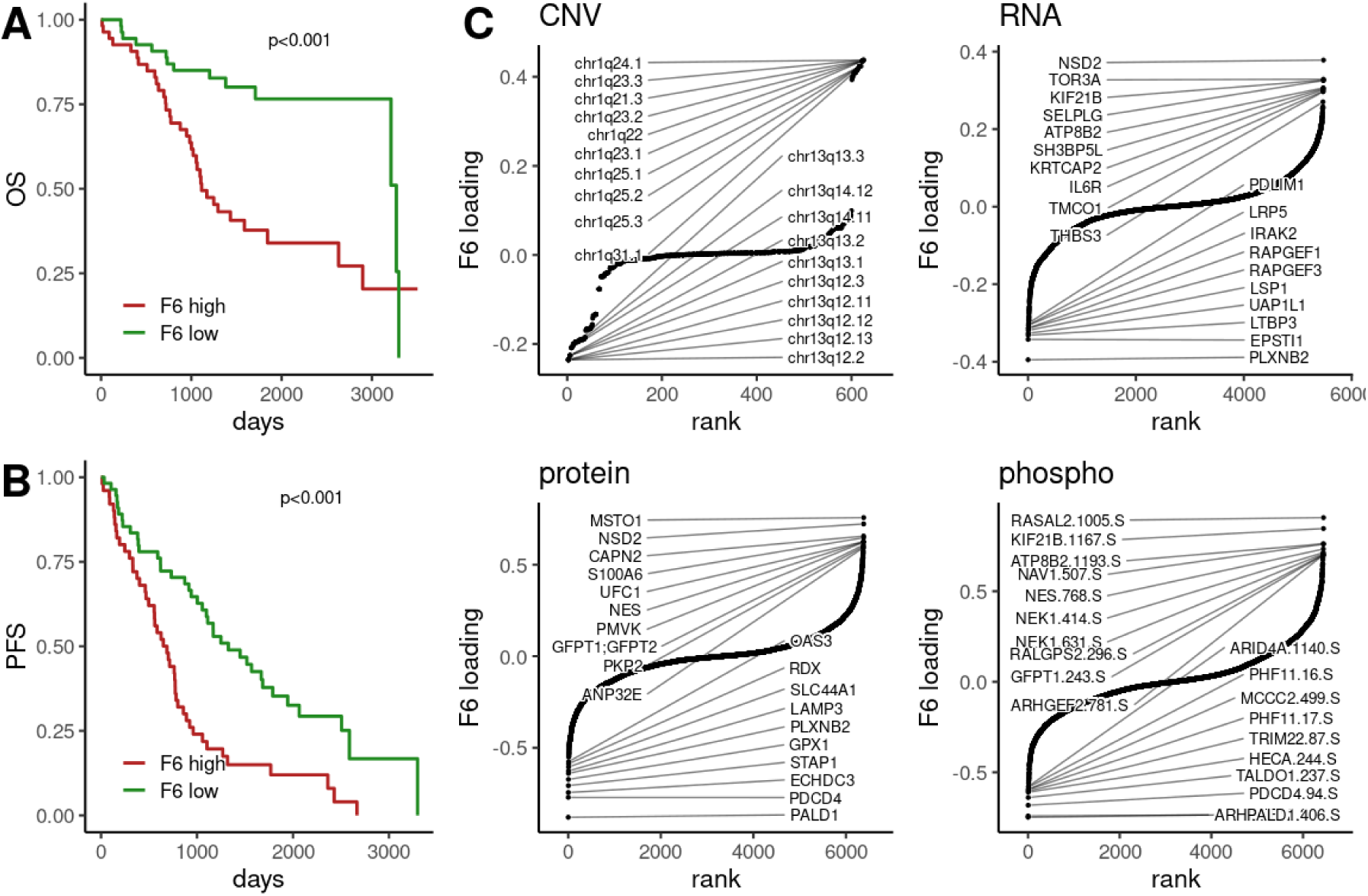
Prognostic value of MOFA2 factor 6. **A+B**: Kaplan-Meier curves for overall survival (OS) (**A**) or progression-free survival (PFS) (**B**) comparing patients with high (>median) or low (<= median) factor 6 score. P-value from log-rank test. **C**: factor 6 loadings in each modality with top features highlighted.

### Uni-modal proxy scores and validation

To enable clinical applications and validate our score in external, single-omics cohorts, we used an elastic net regression approach to derive compact uni-modal proxy scores for F6. Although about 100 features were necessary to obtain R^2^ values of 75% for an RNA proxy, even 10 features were sufficient to get comparable Hazard ratios in a Cox regression (**Figure 5A-C**), and we found similar performance for the protein proxy for F6. We used the same method to derive an RNA proxy for the protein-based risk score, which despite lower R^2^ values reached comparable Hazard ratios even though it required more features for PFS (which the risk score was trained on) (**Figure S5**).

**Figure 5:**
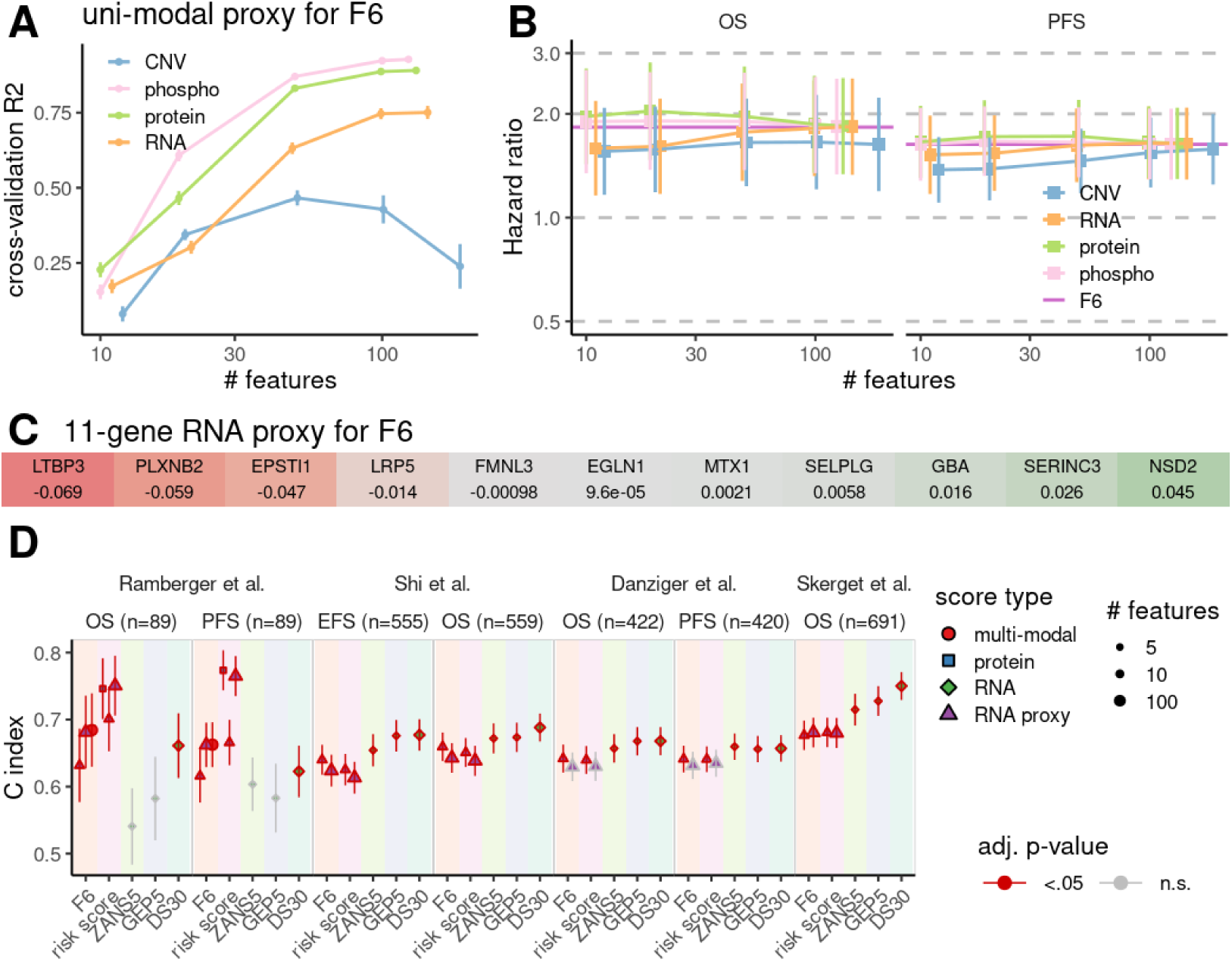
Validating MOFA2 factor 6 and the protein-based risk score using uni-modal proxies. **A**: R^2^ of elastic-net regression as function of the number of active features in uni-modal proxy scores for factor 6 based on CNV, RNA or protein data. Error bars denote standard deviation from 5fold cross-validation. **B**: Cox regression of factor 6 proxy scores with R-ISS as covariate. **C**: Coefficients of an 11-gene RNA proxy for factor 6. **D**: Cox regression analyses (using ISS as covariate) of proxy scores for factor 6 and the protein-based risk score, and the GEP5, ZANS5 and DS30 signatures in multi-omic data of Ramberger et al., microarray data of Shi et al. and Danziger et al. and bulk RNAseq data of Skerget et al.. OS/PFS/EFS: overall/progression-free/event-free survival.

With these RNA-based proxy scores at hand, we went on to validate our findings in three external cohorts: two microarray studies (GSE136337 and GSE24080) (Danziger et al., 2020; Shi et al., 2010), and the CoMMpass RNAseq study (Skerget et al., 2024). We used Cox regression with ISS as covariate to test the F6 score, the protein-based risk score and RNA and protein proxy scores using ∼10 or ∼100 features in the respective modalities of our dataset, as well as the GEP5 (Heuck et al., 2014), ZANS5 (Zamani-Ahmadmahmudi et al., 2021) and DS30 signatures (Bohra et al., 2026). We then compared this to the three validation cohorts, where we evaluated the different RNA proxies for F6 and the risk score together with GEP5, ZANS5 and DS30 and used the non-parametric C-index to compare scores derived from molecular data with different ranges and scales (**Figure 5D**). While the protein-based risk score performs best in our cohort with a C-index of ∼0.75, the performance of its RNA proxies drops to 0.62-0.68 in the validation cohorts. As a result of an unsupervised multi-omic decomposition less prone to overfitting, the F6 score validates somewhat more consistently, with values of 0.67 in our dataset and 0.64 in the other cohorts. Importantly, the 10-feature proxies reach comparable or even better values in the validation datasets than the 100-feature proxies, indicating that compact scores are more robust across cohorts and platforms. The GEP5, ZANS5 and DS30 scores, on the other hand, show C-indices of ∼0.66 in the microarray cohorts (down from > 0.7 in Skerget et al.), and drop to lower values in the smaller Ramberger et al. dataset, where only DS30 still reaches significance.

## Discussion

Our reanalysis of a multi-omics primary multiple myeloma dataset reveals that the functional consequences of genomic drivers are best captured at downstream molecular layers. Supervised classification demonstrated that the proteome and phosphoproteome carry the strongest signal for identifying genomic alterations, outperforming copy-number and transcriptomic data. This hierarchy aligns with a central dogma-like propagation of information, where direct *cis*-acting dosage effects manifest primarily at the RNA level, but the disease-defining molecular state is ultimately encoded in the proteome and post-translational signaling networks. Strikingly, sparse models relying exclusively on *trans*-acting features performed on par with models trained on all features, indicating that the most informative biological readouts are not local genomic effects but rather distal, network-level responses. These findings collectively suggest that the molecular state driving multiple myeloma biology and clinical outcomes is more faithfully captured at the proteomic and phosphoproteomic, rather than genomic level.

At the center of this multi-omic landscape lies Factor 6 (F6), a continuous latent axis that captures a nuanced biological spectrum rather than a binary cytogenetic category. Biologically, F6 balances NSD2-driven epigenetic remodeling at its positive end against immune infiltration at its negative end. Clinically, this axis captures the aggressive, high-risk biology of gain(1q) and del(13q) previously characterized by Skerget et al., but adds critical resolution by explicitly incorporating the tumor immune microenvironment and the effects of recurring translocations. In our dataset, F6 only weakly correlates with previously published prognostic gene expression signatures (GEP5 and ZANS5), or indices associated with proliferation and NF-κB signaling; instead, it reflects a coordinated, immune-modulated state that more accurately stratifies patient prognosis.

One key insight from our results is that the most prognostically relevant biological factors are often not the dominant axes of variance. While principal component analysis readily isolates the largest sources of technical or biological variation, these dominant components frequently lack direct clinical relevance. Instead, F6 emerges as a modest-variance signal that integrates subtle, cross-modal covariance across genomic, transcriptomic, proteomic, and phosphoproteomic layers. As an unsupervised consensus of multi-omic variance, the F6 score may be less prone to overfitting and platform-specific bias than the protein-based risk score by Ramberger et al.. This underscores how multi-omic analysis prioritizes coordinated, cross-layer signals to derive robust and reproducible biomarkers, uncovering prognostic axes that single-omics approaches using supervised, single-cohort optimization would miss.

To translate these complex latent structures into clinically actionable tools, we derived sparse, single-modality proxies. The 11-gene RNA proxy for F6 faithfully recapitulates the multi-omic factor and demonstrates remarkable robustness across different datasets, maintaining prognostic significance in both older microarray cohorts and a contemporary RNAseq dataset. This validates the utility of sparse linear models as a bridge between high-dimensional discovery and routine clinical implementation. By distilling a proteogenomic state into a compact, assayable signature, we provide a pragmatic pathway for risk stratification without requiring expensive, multi-omic profiling in the clinic.

In conclusion, our work demonstrates that trans-acting network responses, rather than isolated cis-acting genomic lesions, are molecularly relevant expressions of multiple myeloma biology and prognosis. The molecular cascade culminates in proteomic and phosphoproteomic states that integrate epigenetic and immune signals into a unified prognostic axis. By leveraging unsupervised multi-omic factor analysis to screen for these subtle, cross-layer signals, and by translating them into sparse, single-modality proxies, we provide a robust and clinically viable framework for risk stratification. This approach not only refines our understanding of myeloma pathogenesis but also offers a clear path toward integrating complex proteogenomic insights into routine clinical decision-making.

## Methods

### Data Preprocessing

We reanalyzed a multi-omics dataset of multiple myeloma comprising whole-genome copy-number variation (CNV) data from nanopore sequencing, transcriptomic (RNA-seq), proteomic (TMT-based), and phosphoproteomic measurements, all derived from the same cohort described in Ramberger et al. Patient annotations, CNV estimates in genomic bins, and normalized expression data for RNA-seq, proteomic, and phosphoproteomic results were extracted from Supplementary Tables 2–7 of the original publication. Only patients with complete data across all four modalities were retained for downstream analysis. Features (loci, genes, proteins, or phosphosites) with missing values or lacking gene annotation were excluded, yielding a final cohort of 120 patients with 626 CNV loci, 5477 genes, 6383 proteins, and 6446 phosphosites.

Clinical and genomic annotations were integrated from the original study and supplemented with additional data including plasmacytosis (with and without MACS sorting), genetic subgroups, chromosomal aberrations (del 9q, gain 1q, del 13q, del 17p, t(4;14), t(11;14), t(14;16)), and ISS/R-ISS staging. RNA quality (DV200) was incorporated as a covariate. The protein-based risk score was computed using a weighted sum of eight proteins (ANKZF1, UBE2Q1, PDSS2, TFB2M, NCKAP1, PTK2, RDX, BAG4). Additional composite scores were calculated: a proliferation index from the mean expression of 12 cell-cycle genes (Bergsagel et al., 2005), an NF-κB index from the mean expression of 11 NF-κB pathway genes (Ang et al., 2024), a 5-gene version of the GEP-70 score (Heuck et al., 2014), a ZANS5 score (Zamani-Ahmadmahmudi et al., 2021), and the DS30 signature (Bohra et al., 2026).

### Batch Correction and Principal Component Analysis

To account for technical batch effects, we applied the removeBatchEffect function from the limma R package. RNA-seq data were corrected for both MACS protein content and RNA quality (DV200), whereas protein and phosphoproteomic data were corrected for MACS protein content alone. Principal component analyses (PCAs) were performed on all four modalities before and after batch correction to assess data quality and batch effect removal. Silhouette widths were computed to evaluate the separation of patients by genomic events (HRD, gain(1q), gain(9q), del(13q), del(17p)) in PC space.

### Supervised Classification with (s)PLS-DA

We assessed the contribution of each omics modality to the classification of key genomic events using partial least squares discriminant analysis (PLS-DA) and sparse PLS-DA (sPLS-DA) implemented in the mixOmics R package (Rohart et al., 2017). For each genomic event (HRD, gain(1q), gain(9q), del(13q), del(17p), t(4;14), t(11;14), t(14;16)), we trained separate PLS-DA and sPLS-DA models on each modality independently, using two latent components. For sPLS-DA, we used 100 features for the first and 50 for the second component. Two sPLS-DA strategies were compared: (i) using all features, and (ii) excluding features located on the target locus (i.e., restricting to *trans*-acting features only) to assess whether models relied on direct on-target signals.

Model performance was evaluated using 50 repetitions of 5-fold cross-validation (3-fold for events with fewer than 20 positive cases), with area under the receiver operating characteristic curve (AUROC) as the primary metric. The squared loadings of each feature were extracted to assess the contribution of *cis* (on-target) versus *trans* (off-target) features. Wilcoxon rank-sum tests compared *cis* and *trans* loading distributions, and bootstrap confidence intervals (500 resamples) were computed for the difference in median loadings between *cis* and *trans* features.

### Gene Set Enrichment Analysis

To identify biological pathways associated with genomic events, we performed gene set enrichment analysis on the ranked features from the dense PLS-DA models using the tmod R package (Zyla et al., 2019). Gene sets were obtained from the Molecular Signatures Database (MSigDB), including Hallmark and Reactome gene sets. Features were ranked by the sum of squared loadings across components, and the CERNO test was applied. Pathways were considered significantly enriched at an adjusted *P*-value < 0.001 and AUC > 0.65, and only pathways significant in at least two comparisons were retained for downstream analysis. Enrichment effect sizes (AUC) were clustered using hierarchical clustering with Ward’s method.

### Multi-Omics Factor Analysis (MOFA2)

To capture shared and modality-specific latent structure across all four omics layers, we performed Multi-Omics Factor Analysis using MOFA2 (v0.7.3) (Argelaguet et al., 2020) with 10 latent factors. The explained variance per factor and modality was computed, and the optimal number of factors was evaluated by examining cumulative variance explained and factor stability across factor numbers (2–50).

Factor scores were associated with clinical and molecular variables using non-parametric tests: Wilcoxon rank-sum for binary variables, Kruskal–Wallis for multi-class variables, and Spearman correlation for continuous variables. *P*-values were adjusted using the Benjamini-Hochberg procedure. Gene set enrichment was performed on factor loadings using the same CERNO framework as for PLS-DA.

### Survival Analysis

Prognostic associations were assessed using Cox proportional hazards models adjusted for R-ISS stage where available or ISS otherwise, using standardized factor scores. Univariate hazard ratios (HR), 95% confidence intervals, and *P*-values were computed for progression-free survival (PFS) and overall survival (OS).

### Uni-modal Proxies for Latent Factors

To identify interpretable, single-modality surrogates for the multi-modal latent factors (particularly Factor 6, which showed the strongest prognostic signal), we trained sparse linear models using glmnet (elastic net, α = 0.5). For each modality (CNV, RNA, protein), we fitted regularized regression models predicting Factor 6 scores on standardized modality data, scanning feature counts from 10 to 200. Cross-validated *R*^2^ was computed across 20 repeats of 5-fold CV. The optimal lambda for each feature count was selected from the full-data fit, and out-of-sample performance was evaluated on held-out folds.

### External Validation

Uni-modal proxies for Factor 6 and the protein risk score were validated in three independent multiple myeloma cohorts:

1. CoMMpass (Skerget et al., 2024): RNA-seq data from primary bone marrow samples with clinical annotations including ISS stage and overall survival.
2. GSE136337 (Danziger et al., 2020): Microarray-based transcriptomic data with PFS and OS.
3. GSE24080 (Shi et al., 2010): Microarray-based transcriptomic data with EFS and OS.

For each validation cohort, proxy scores were computed as weighted sums of the selected standardized features using coefficients from the training cohort. Cox models adjusted for ISS stage were fitted to assess prognostic concordance. A summary table of hazard ratios across all cohorts, endpoints, and score types was compiled for comparative assessment.

### Software and Reproducibility

All analyses were performed in R (v4.3.2) using packages mixOmics (6.26.0), limma (3.58.1), glmnet (v4.1-8), tmod (v0.50.13), msigdbr (v7.5.1), survival (v3.8-3) and survminer (v0.5.0), while the MOFA2 analysis was run in python (v3.12) using mofapy2 (v0.7.3). Code is available at [github.com/bihealth/multiple-myeloma-multiomics].

## Acknowledgements

This work was supported by the Federal Ministry of Research, Technology and Space (BMFTR), as part of the National Research Cores for Mass Spectrometry in Systems Medicine under grant agreement no. 03LW0239K (MSTARS).

## AI usage

Generative AI tools ChatGPT, Claude and Qwen3.5 were used during the preparation of this manuscript to support literature review, brainstorming, creation of scripts to generate figures, and the drafting and editing of parts of the manuscript. All analysis, interpretation, and conclusions are the authors’ own.

## Supplementary Figures

**Figure S1:**
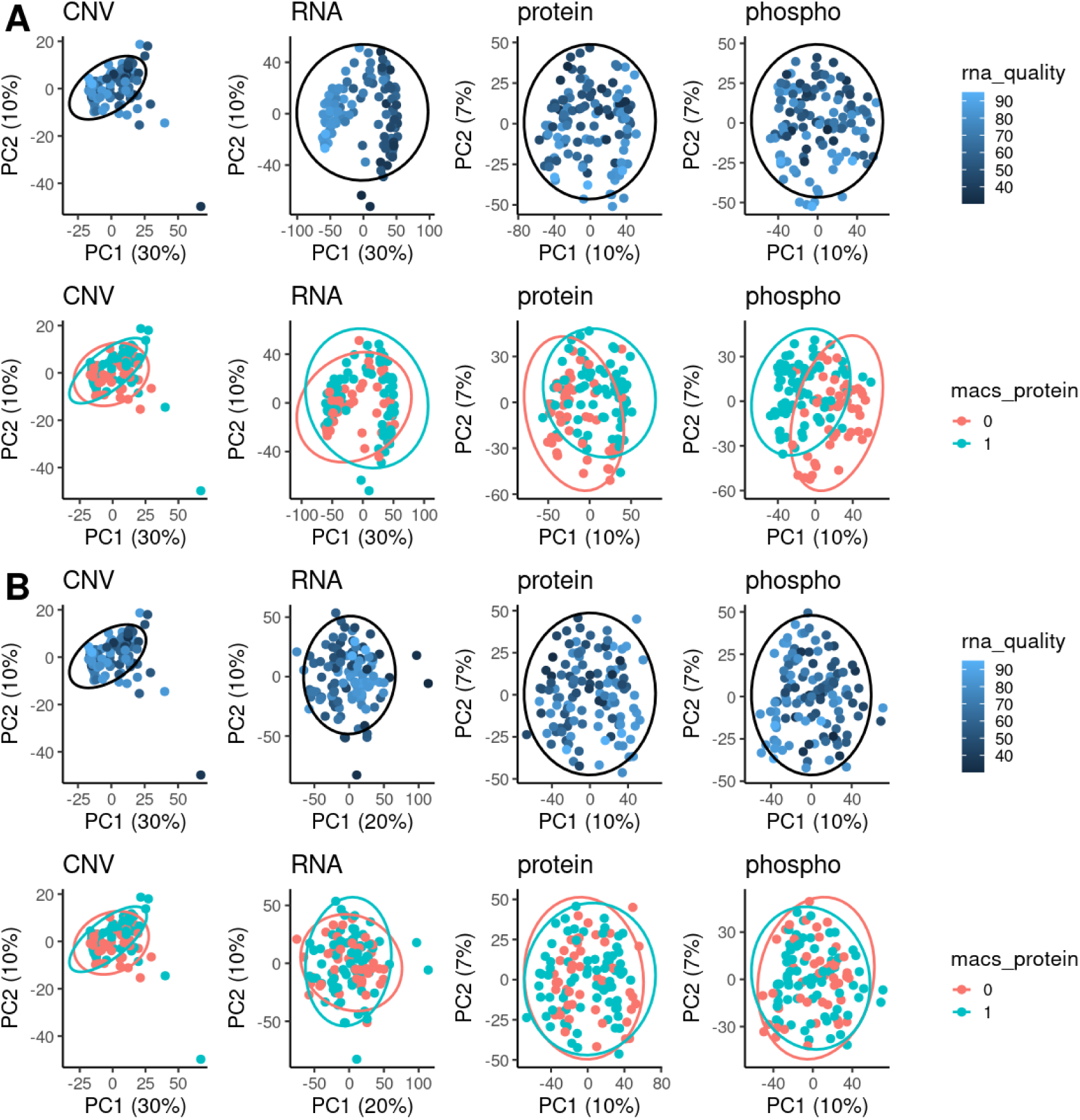
Principal component analyses per modality colored by RNA quality (top row) or MACS sorting (bottom row) before (**A**) and after (**B**) batch correction with LIMMA.

**Figure S2:**
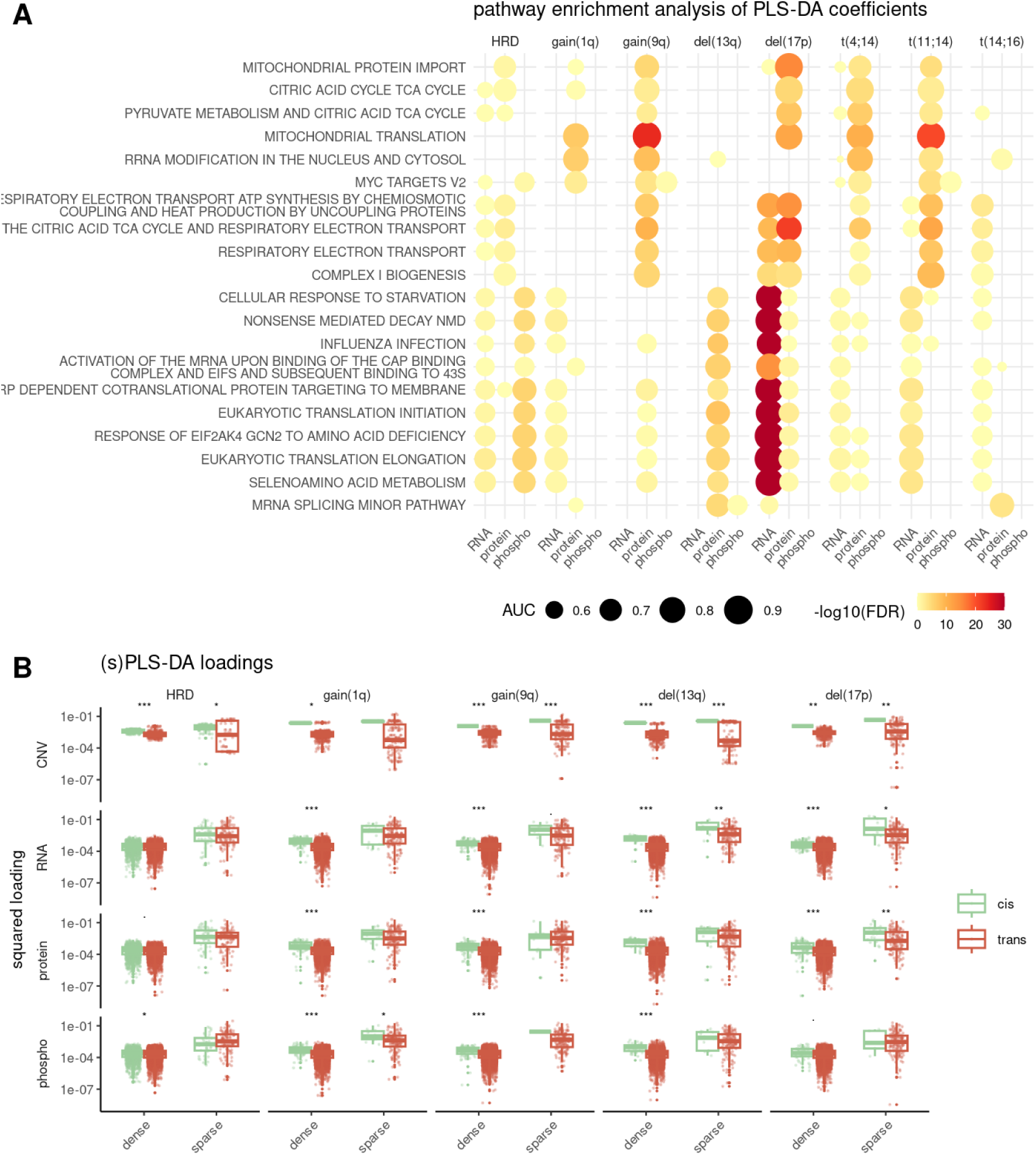
**A**: Gene set enrichment analysis of PLS-DA loadings per modality and genomic event. Symbol size indicates effect size (area-under-curve), color scale indicates adjusted p-value. **B**: Comparing loadings for cis and trans features in dense or sparse PLS-DA analyses. P-values from Wilcoxon test. ***: p < 0.001, **: p < 0.01, *: p < 0.05.

**Figure S3:**
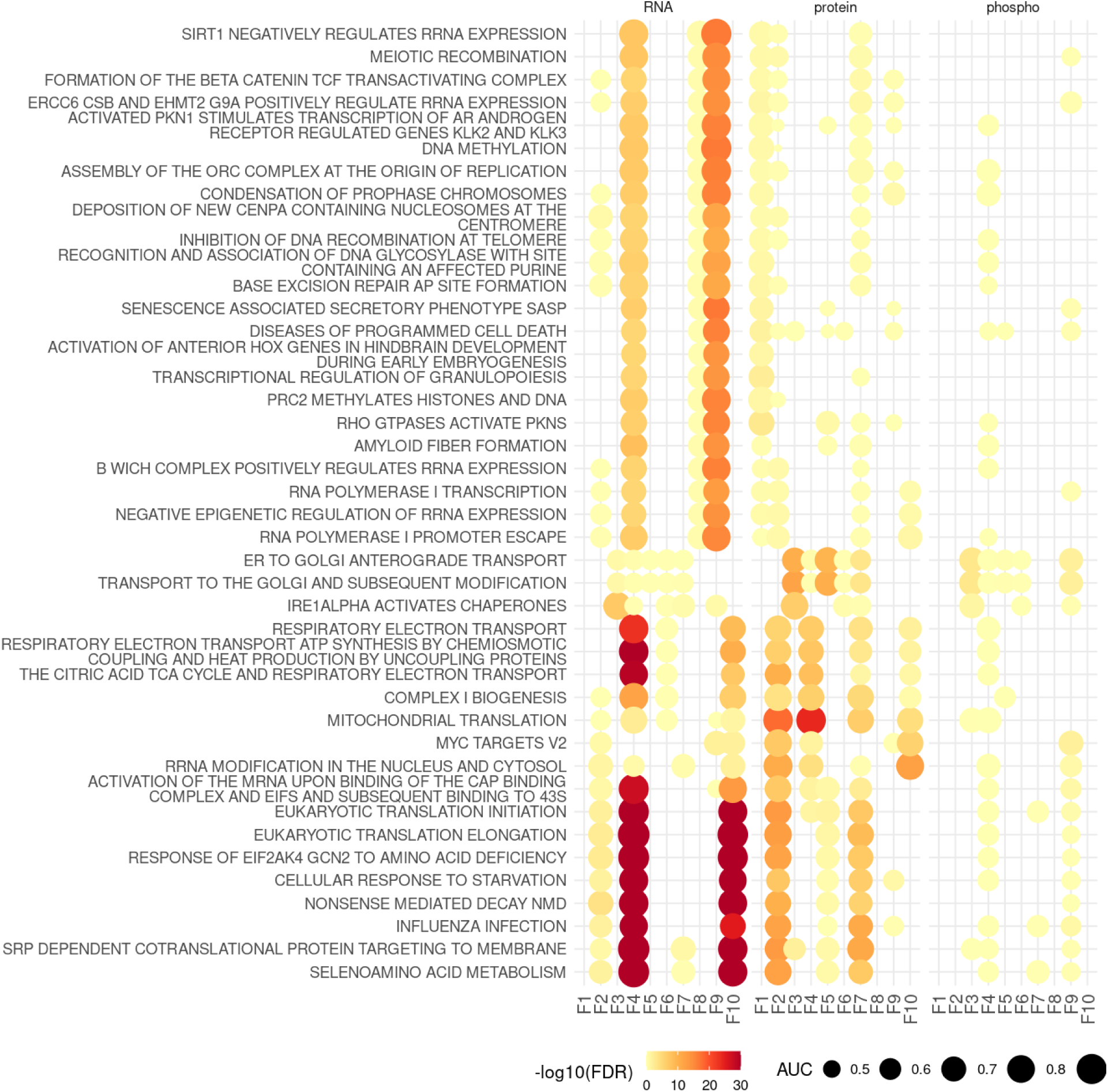
Gene set enrichment analysis on MOFA2 factors using REACTOME and HALLMARK gene sets. Symbol size indicates effect size (area-under-curve), color scale indicates adjusted p-value.

**Figure S4:**
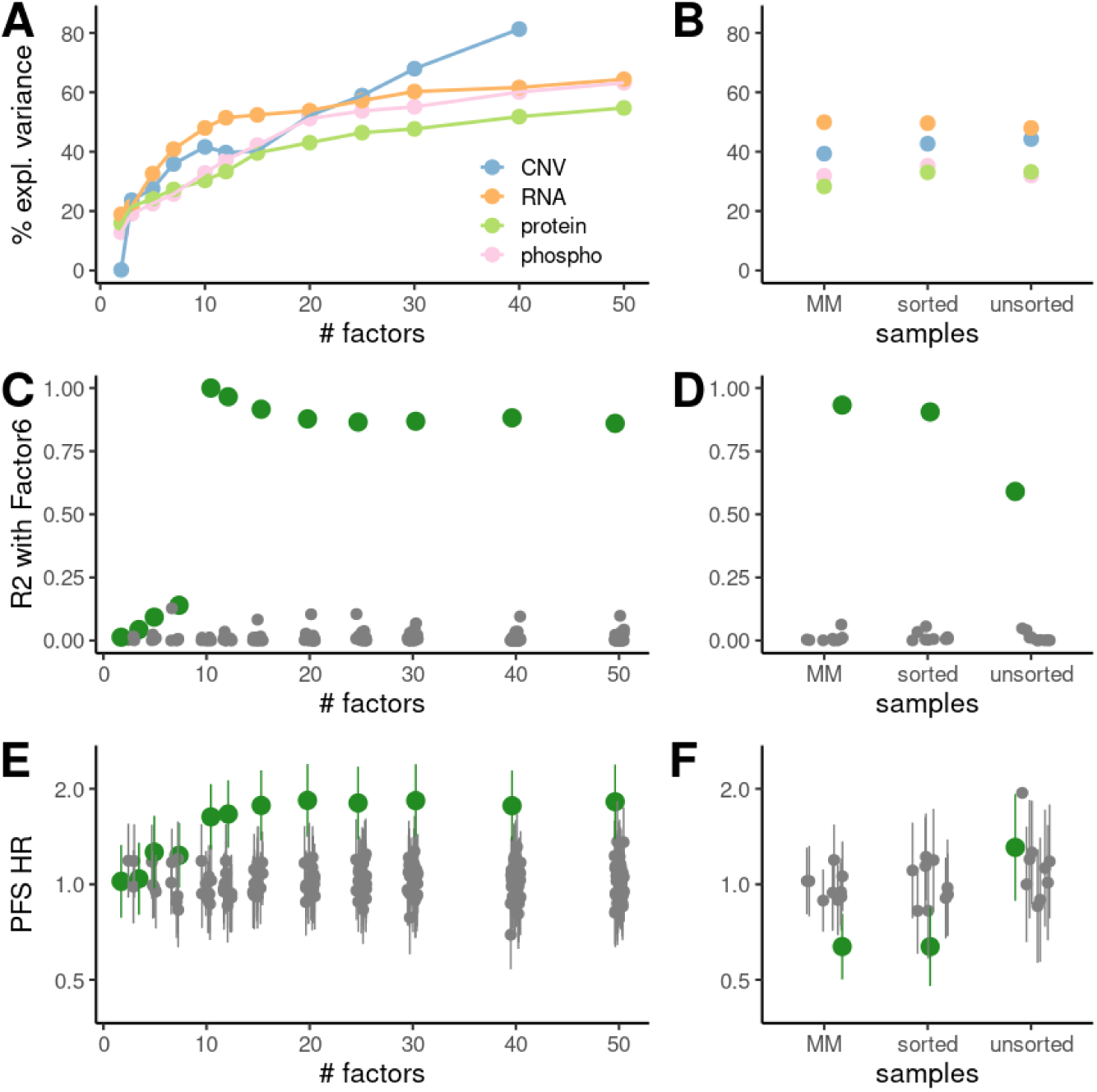
Running MOFA2 models with different numbers of factors (left) or on different subsets of samples (right; MM: only multiple myeloma patients, sorted/unsorted: CD138+ sorted or unsorted input) **A+B**: Total variance explained per modality. **C+D**: R^2^ with MOFA2 factor 6 from Fig. 3. Best scoring factor is highlighted. **E+F**: Cox regression analyses for all factors of these models. Best-scoring factor from C or D is highlighted.

**Figure S5:**
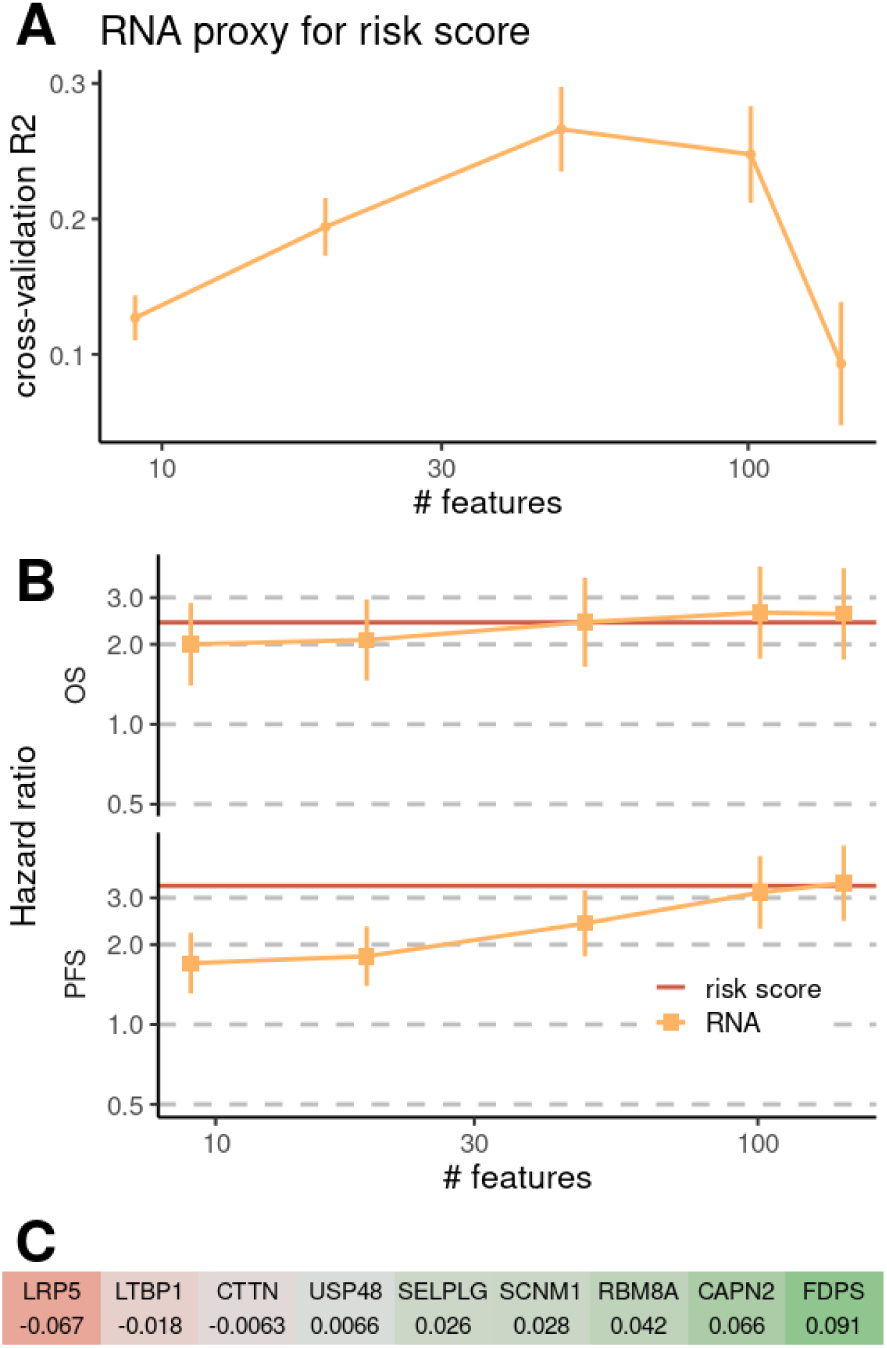
**A**: uni-modal proxy score (based on RNA) for the protein-based risk score of Ramberger et al. **B**: Cox regression for this uni-modal proxy. **C**: coefficients of a 9-gene RNA proxy.

